# Control of Shoot Meristem Size by an Aminoacyl tRNA Synthetase, *OKI1*, in *Arabidopsis*

**DOI:** 10.1101/548164

**Authors:** Munenori Kitagawa, Rachappa Balkunde, Huyen Bui, David Jackson

## Abstract

In plants, the stem cells that form the shoot system reside within the shoot apical meristem (SAM), which is regulated by feedback signaling between the WUSCHEL (WUS) homeobox protein and CLAVATA (CLV) peptides and receptors. WUS-CLV feedback signaling can be modulated by various endogenous or exogenous factors such as chromatin state, hormone signaling, reactive oxygen species (ROS) signaling and nutrition, leading to a dynamic control of SAM size corresponding to meristem activity. Despite these insights, however, the knowledge of genes that control stem cell fate is still limited, and in particular the regulation by ROS signaling is only beginning to be comprehended. Here, we report a new regulator of SAM size, *OKINA KUKI* (*OKI1*), which is expressed in the SAM and encodes a mitochondrial aspartyl tRNA synthetase (AspRS). *oki1* mutants display enlarged SAMs with abnormal expression of WUS and CLV3, and overaccumulation of ROS in the meristem. Our findings support the importance of normal AspRS function in the maintenance of the WUS-CLV3 feedback loop and SAM size.

## Introduction

Stem cells are responsible for the generation of new tissues and organs in multicellular organisms, while maintaining themselves as pluripotent initials. In plants, stem cells reside in meristems, and the shoot apical meristem (SAM) can generate all shoot organs, such as leaves and flowers. Semi-permanent stem cells are found at the apical region of the SAM, in the central zone (CZ), and the organizing center (OC) below is a group of niche cells that provides cues to the CZ for stem cell maintenance. The daughter cells produced by stem cell divisions are displaced to the peripheral zone (PZ), where they form organ primordia, and in turn differentiate into lateral organs. Thus, stem cell fate and differentiation are precisely regulated depending on position within the SAM, allowing maintenance of the stem cell population and meristem size.

For such position dependent maintenance of the stem cell niche, plants have developed a negative feedback signaling pathway. The homeodomain transcription factor WUSCHEL (WUS) is expressed in the OC, and moves through plasmodesmata into the CZ to activate the expression of its negative regulator CLAVATA3 (CLV3) (Daum et al. 2014; Fletcher et al. 1999; Mayer et al. 1998; Perales et al. 2016; Yadav et al. 2011). CLV3 is a secreted peptide perceived by the leucine-rich repeat receptor like protein kinases (LRR-RLKs) CLAVATA 1 (CLV1) and the related BARELY ANY MERISTEM (BAMs) as well as by the LRR receptor like protein CLAVATA2 (CLV2) in a complex with the CORYNE (CRN) pseudokinase, which together repress the expression of WUS in the OC (Clark et al. 1997; DeYoung et al. 2006; DeYoung and Clark 2008; Hu et al. 2018; Kayes and Clark 1998; Miwa et al. 2008; Muller et al. 2008; Nimchuk et al. 2011; Nimchuk et al. 2015; Ogawa et al. 2008; Shinohara and Matsubayashi 2015). This WUS-CLV3 negative feedback loop establishes a self-correcting mechanism that maintains proper size of the stem cell pool and the meristem (Brand et al. 2000; Schoof et al. 2000; Somssich et al. 2016).

The WUS-CLV negative feedback loop is also regulated by various endogenous and exogenous signals. For example, precise WUS expression patterns require chromatin regulators, and mutants of these factors display bigger and disorganized meristems (Graf et al. 2010; Kaya et al. 2001). In addition, cytokinin promotes WUS expression and in turn facilitates the proliferation of stem cells, leading to an increase in SAM size (Chickarmane et al. 2012; Gordon et al. 2009; Gruel et al. 2016; Landrein et al. 2014). Cytokinin signaling also controls SAM size depending on the availability of nutrients in a WUS-dependent manner, allowing plants to optimize shoot organogenesis according to resource availability (Landrein et al. 2018). By contrast, auxin signaling negatively regulates the stem cell population by modulating WUS-CLV3 feedback loop through interaction with cytokinin signaling (Shi et al. 2018; Zhao et al. 2010). ROS signaling is also an important regulator of the stem cell population and SAM size. Mutants of a mitochondrial protease and plastid ion channels display abnormal accumulation of ROS at the shoot apices under abiotic stresses, leading to premature termination of the SAM and abnormal growth of calluses at the shoot apices, respectively (Dolzblasz et al. 2016; Wilson et al. 2016). Furthermore, Zeng et al. (2017) reported that superoxide anions are enriched in the stem cells of the SAM and promote WUS expression, whereas hydrogen peroxide accumulates in the PZ to promote differentiation. These findings suggest that the proper accumulation and precise distribution of ROS are crucial for the maintenance of stem cell niches and SAM size. The same applies for regulating the root apical meristem (Jiang et al. 2003; Kong et al. 2018; Tsukagoshi et al. 2010; Yang et al. 2014; Yu et al. 2016; Yu et al. 2013). For example, a recent study revealed that prohibitin (PHB3) regulates ROS homeostasis in roots, and in turn maintains root meristem size and stem cell niches through the functions of its downstream ROS-responsible factors; *phb3* mutants show overaccumulation of ROS in roots, reduced root meristem size and defective quiescent center (Kong et al. 2018).

Despite these insights, the knowledge of genes affecting SAM fate is limited, and how ROS is regulated in the SAM is only beginning to be understood. In this study, we identify the *OKINA KUKI* (*OKI1*, Japanese for big stem) gene as a new regulator of SAM size. *oki1* mutant seedlings have enlarged SAMs and abnormal expansion of WUS and CLV3 expression. *OKI1* encodes a mitochondrial AspRS that is expressed in the SAM, and the mutant meristems show abnormal accumulation of ROS, suggesting a possible mechanism for abnormal meristem development. Collectively, our discoveries suggest that normal function of the AspRS OKI1 is required to maintain the WUS-CLV3 feedback loop and SAM size.

## Results

In an ethyl methyl sulfonate (EMS) screen for new mutants affecting shoot development in *Arabidopsis*, we identified a mutant with small seedlings and the first 4-5 leaves were narrow and strap shaped (Fig. 1A, B). Later in development, the plants recovered their growth somewhat, and made inflorescence shoots that were often highly fasciated (54%, n= 20/37, Fig. 1D-F). As this phenotype is usually associated with enlargement in meristem size (Fig. 1C), we measured shoot meristems from the mutants and their normal siblings (Fig. 1G-J). The mutant meristems were normal at 8 days after planting (DAP), but were significantly wider and taller than their siblings at 12 DAP (N = 10-15; P < 0.01, Tukey HSD test). To understand the cellular basis of this phenotype, we fixed and sectioned both shoot and root apices for imaging in the confocal microscope. Normal shoot meristems have cells arranged in two regular outer layers, the L1 and L2, with an inner group of L3 cells (Fig. 1K, L). In contrast, the mutant shoot meristems had highly disorganized cell arrangement, and the regular L1 and L2 layer structure was less evident (Fig. 1M, N). Similar phenotypes were found in root meristems, where cells are again normally arranged in regular radial layers (Fig. 1O, P). In the mutants, we again saw evidence of irregular layers, with cells expanding into the space of the adjacent layers, and irregular planes of cell division (Fig. 1Q, R). Because of the prominent fasciated stem phenotype, we named this mutant *okina kuki* (*oki1*, Japanese for big stem).

**Figure 1.**
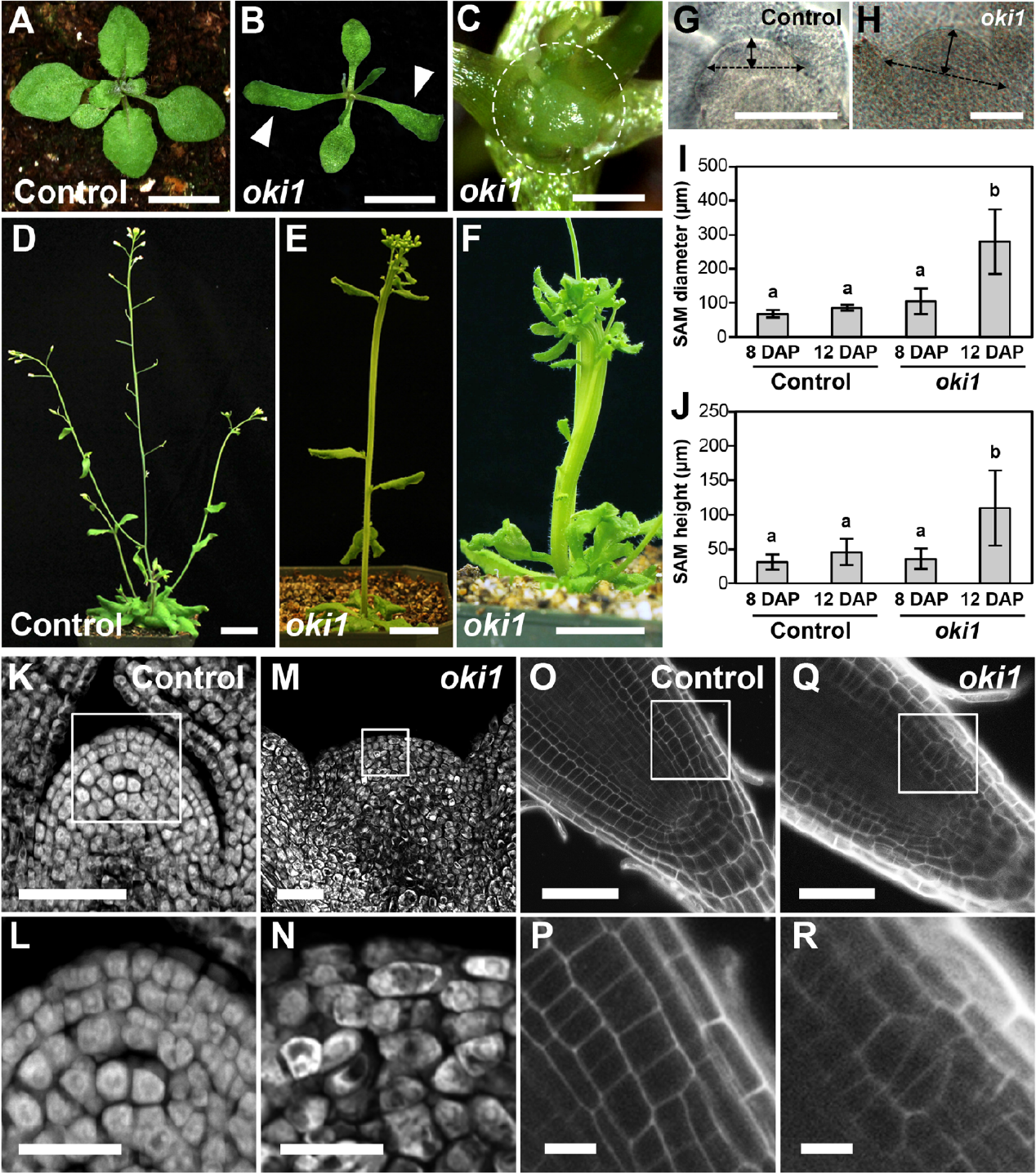
Shoot and root meristems are affected in *oki1* mutants. (A, B) *Oki1* displayed delayed growth and narrower true leaves (B, arrowhead) in seedlings compared with control line (Col-0, A). Scale bars = 5 mm. (C) *Oki1* mutants have enlarged SAM (dashed circle). Scale bar = 500 μm. (D-F) The inflorescence stems of control line (D) and *oki1* (E, F); *oki1* was often fasciated (E and F). Scale bars 22 cm (D) and 1 cm (E, F). (G, H) Cleared SAM from the control line (G) and *oki1* (H) at 12 DAP. Solid and dashed double-headed lines display the SAM height and diameter, respectively. Scale bar 100 = μm. (l, J) The SAM Of *Oki1* was significantly larger at 12 DAP. N = 10-15. Bars topped by different letters are significantly different at P < 0.01 (Tukey HSD test). Cell arrangements were disorganized in the *oki1* SAM and root tips. Confocal images of Eosin Y-stained SAM sections of control line (K) and *oki1* (M), and PI-stained root tips Of control line (O) and *Oki1* (Q). L, N, P, R show magnified images Of the boxed regions in K, M, O and Q respectively. Scale bar = 50 μm (K, M, O, Q), 20 μm (L, N) and 10 μm (P, R).

The *oki1* mutant was identified in Columbia-0 (Col-0), so to identify the underlying gene we crossed it to the *Landsberg erecta* (Ler) ecotype, and made a bulk mutant pool from the F2 mapping population. We next used whole genome sequencing, followed by analysis using the SHOREmap pipeline (Schneeberger et al. 2009) to map the mutation to chromosome 4, between ~ 15-17 Mb. Further fine mapping, combined with analysis of gene sequences within the mapping interval revealed a candidate mutation in AT4G33760, a gene encoding an aspartyl tRNA synthetase. A single base pair change, G347A, in the first exon of this gene led to a single predicted amino acid change, G116D, in the anti-codon binding domain of this protein (Fig. 2A, B and Fig. S1). To confirm this was the correct mutation underlying the phenotype, we obtained a T-DNA insertion, SAIL_358_B08, in the 10^th^ exon of AT4G33760, and when plants heterozygous for this insertion were crossed to *oki1* plants, the T-DNA insertion allele failed to complement the *oki1* phenotype, as the progeny segregated ~ half with *oki1* phenotype (Fig. 2C and Fig. S2A), and these plants were confirmed as being heterozygous for the *oki1* EMS allele and the T-DNA mutations. Furthermore, we were able to complement the *oki1* mutation using a TAC clone, JAtY59F05, containing At4g33760 (Fig. S2B), but not when At4g33760 was mutated in this TAC, together indicating that the gene underlying the *oki1* mutation was correctly identified.

**Figure 2.**
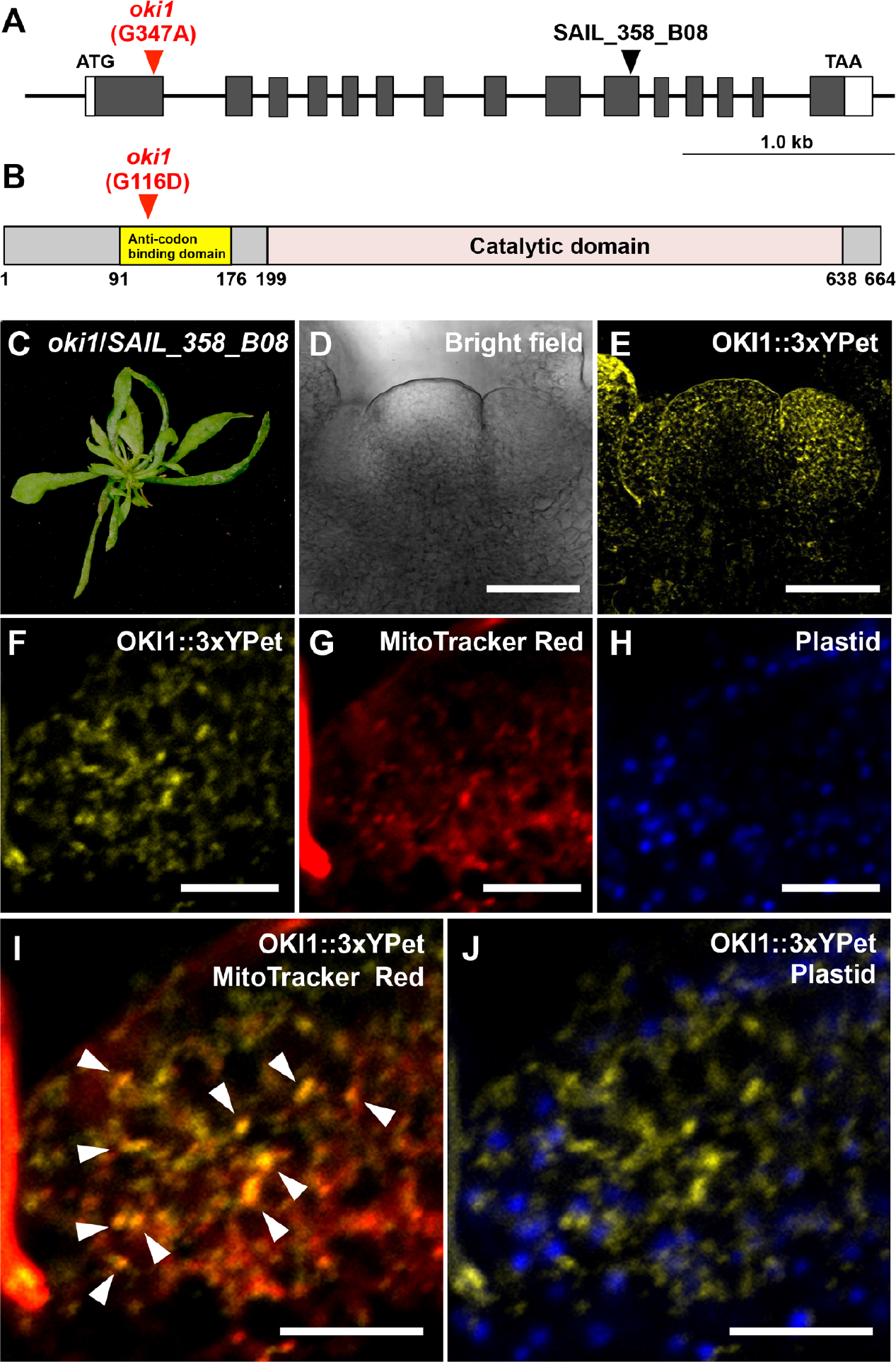
The causal gene of *oki1* encodes a mitochondrial aspartyl tRNA synthetase (AspRS). (A) Diagram of the intron-exon structure of *OKI1* gene (*At4g33760*). UTRs are indicated by white boxes, coding regions Of exons by grey boxes. Solid lines indicate the intergenic regions and introns. The nucleotide substitution in *oki1* (G347A) is shown in red arrowhead. T-DNA insertion site for SAIL_358_B08 is shown in black arrowhead. (B) Schematic diagram of the domain structure of OKI1 protein. Predicted anti-codon binding domain (91-176 aa) and the catalytic domain of AspRS (199-638 aa) are shown in yellow and pink boxes, respectively. Glycine 116 within the anti-codon binding domain was changed to aspartic acid in *oki1* mutants (red arrowhead). (C) F1 plants from crosses between *oki1* and *SAIL_358_808* showed enhanced *oki1* narrow leaf phenotype (alsosee Fig. S2A). (D, E) Bright field and fluorescence images of the inflorescence SAM showing OKI1∷3xYPet driven under its native promoter in *oki1* background. pOK11-OK11∷3xYPet is expressed in the meristem and shoot apex. Scale bar = 50 μm. (F-H) Subcellular localization Of (F), mitochondria stained with MitoTracker Red (G), autofluorescence of plastids (blue) (H) in the cells Of the SAM. Merged images of F with G (l) and F with H (J) are shown. OKI1∷3xYPet localized to mitochondria (overlap shown by white arrows), but not plastids (blue) in the meristem. Scale bar = 10 μm.

Aminoacyl tRNA synthetases play a critical role in cellular metabolism by charging tRNAs with their cognate amino acid for protein synthesis. *Arabidopsis* encodes three aspartyl tRNA synthetases, and the one encoded by AT4G33760 is expressed ubiquitously, with strongest expression in seedling leaves and vegetative shoot meristems (Fig. S3). The product of AT4G33760 is predicted to localize to mitochondria and/or chloroplasts (Duchêne et al. 2005), while the two other aspartyl tRNA synthetases, At4G26870 and At4G31180 are predicted to encode cytoplasmic proteins (Fig. S4) (Luna et al. 2014). To ask where the OKI1 protein localized, we made a triple YPet fluorescent protein fusion tagged at the C terminus of the predicted coding sequence, in the context of its native regulatory sequences in the TAC clone. This construct was able to fully complement the mutant phenotype, and in mature leaf tissues, we observed OKI1-YPet localization in a punctate pattern that overlapped with a mitochondrial stain, MitoTracker Red (Fig. S5A-D). In shoot meristems, we also saw punctate staining (Fig. 2D, E), that again co-localized with MitoTracker Red, and not with plastids, visualized by autofluorescence (Fig. 2F-J). In summary, we identified *oki1* as a point mutation in an aspartyl tRNA synthetase that localizes to mitochondria in the leaf and shoot meristem, and is a weak allele, since a putative null allele was lethal.

We next ask how the *oki1* mutant interacted with the canonical CLV-WUS feedback pathway that maintains the stem cell population in the shoot meristem. We crossed the mutant to a line carrying a GFP reporter for CLV3, as well as a RFP reporter for WUS expression. In wild type (WT) siblings, these reported the expected expression, with CLV3 expressed in 2 to 3 cell layers of stem cells in an arc at the top of the meristem, and WUS in a cluster of organizing center cells below in red (Fig. 3B, F). In the mutants, the separation of the CLV and WUS domains was maintained, however the meristems were enlarged and irregular, as already described, and the CLV3 and WUS expression domains were expanded (Fig. 3D, H and Fig. S6). We also asked how these mutations interact in double mutant combinations. *wus* mutants make irregular shoots that terminate prematurely, and even after bolting make few, irregular flowers (Fig. 3K, L). In double *wus oki1* mutants, *wus* behaved epistatically, as the double mutants were indistinguishable from *wus* (Fig. 3K-N). *clv3 oki1* double mutants similarly resembled the *clv3* single mutants (Fig. 3O, P), and quantification of phenotypes by measurements of stem thickness indicated that there was no significant difference in stem thickness between *oki1* or *clv3* single mutants and *oki1 clv3* double mutants (Fig.S7). Together these double mutant analyses indicate that *wus* and *clv3* are epistatic to *oki1* in meristem size control.

**Figure 3.**
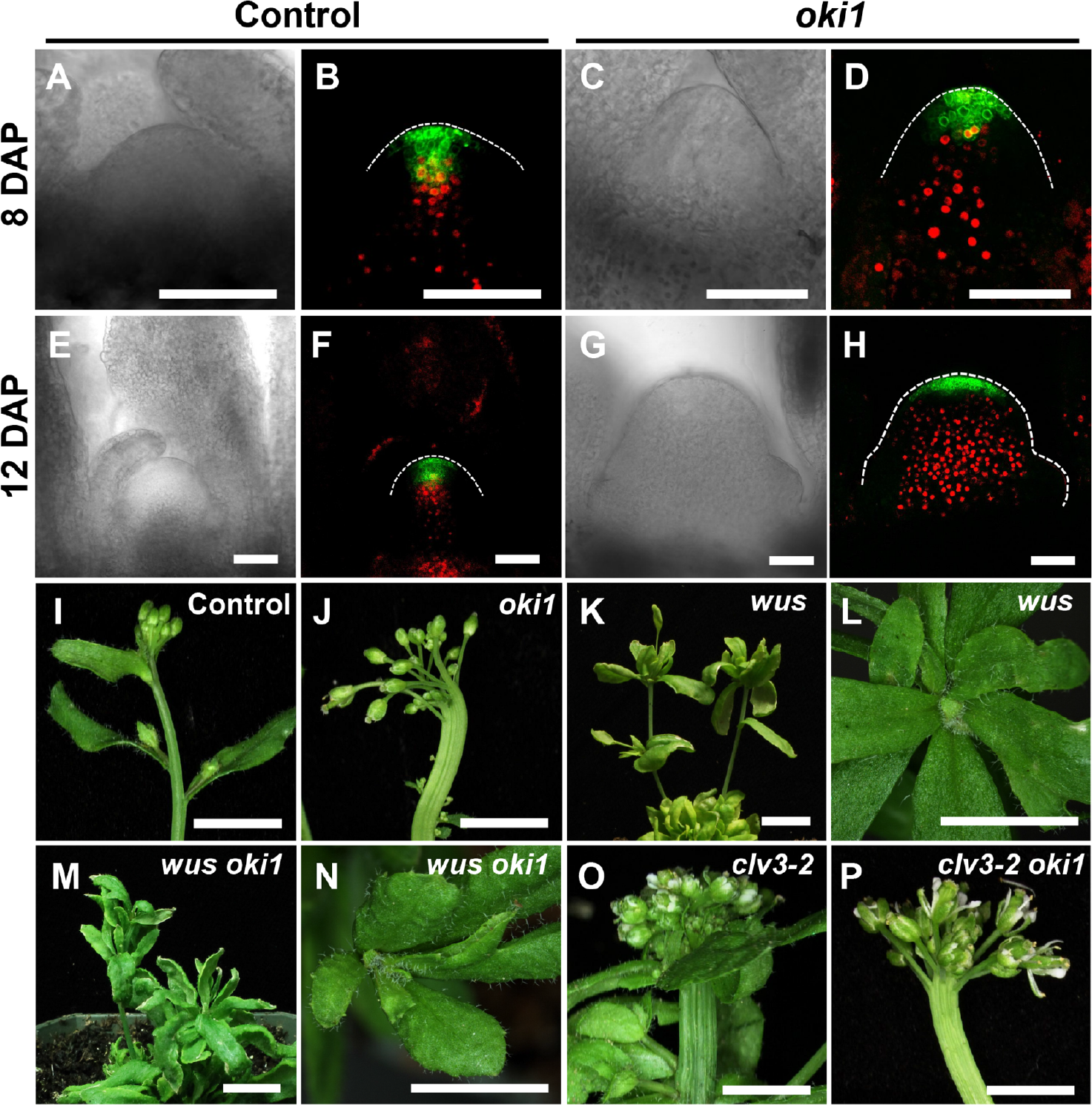
Relationship Of *OKI1* with known meristem signals. (A-H) Confocal images of pCLV3∷mGFP5-ER (green) and pWUS∷dsRED-N7 (red) in the SAM from 8 DAP (A-D) and 12 DAP (E-H) seedlings Of control line (A, B, E, F) and *oki1* (C, D, G, H). Dashed lines display the outlines of the SAM. WUS and CLV3 promoter activities were enlarged in the SAM of *oki1* at 12 DAP. Scale bar = 100 μm. (l, J, K, M, O, P). *wus* and *clv3* are epistatic to *oki1*. Shoot apices Of 35 DAP plants Of control line (I), *Oki1* (J), *wus* (K, L), *wus oki1* (M, N), *clv3-2* (0) and *clv3-2 oki1* (P). Scale bars = 5 mm (l, J, L, N, O, P), lcm (K, M).

Finally, to address the possible mechanism by which *oki1* mutants cause meristem disruption, we reasoned that a block in mitochondrial function by partial loss of an essential translation factor might lead to redox inbalance, which is known to impact meristem size (Zeng et al. 2017). We therefore stained meristems with redox dyes, and indeed found that superoxide and peroxide were upregulated in *oki1* meristems (Fig. 4), suggesting that redox imbalance may cause the increases in meristem size in *oki1*.

**Figure 4.**
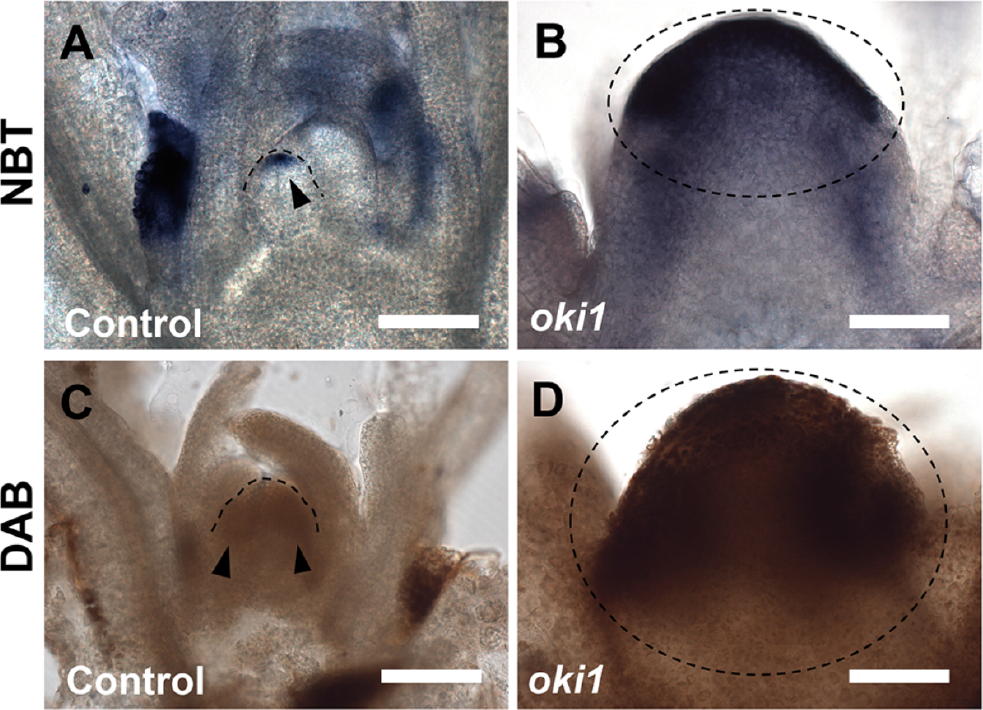
*oki1* overaccumulates superoxide and hydrogen peroxide in the SAM. (A, B) Nitroblue tetrazolium (NBT) staining showed that superoxide is higher in the SAM of *oki1* (B, dark blue stained region within dashed circle) compared with control line (A, blue region shown by arrowhead). (C, D) 3,3’-diaminobenzidine (DAB) staining indicated that hydrogen peroxide levels were higher in the SAM of *oki1* (D, brown and black regions within dashed circle) compared with control line (C, brown regions indicated by arrowheads). Dashed lines in A and C show the outlines of the SAM of the control line. Scale bars = 100 μm.

## Discussion

Meristems are highly ordered structures that initiate new organs throughout the lifecycle, to enable plant survival. Several mutants with disorganized meristems has been described, and among the ones with larger meristems the best understood are the CLV pathway genes, *CLV1*, *CLV2* and *CLV3*, which repress *WUS* to balance the loss of cells from the meristem due to organ initiation with the production of cells by stem cell divisions. This balance is crucial to maintain meristem size, and other genes acting in the peripheral zone of the meristem, either providing feedback to the stem cell niche or promoting the transition of cells into organ primordia, also lead to bigger meristems when mutated (Chuck et al. 2014; Pautler et al. 2015). Hormonal feedback signaling, most notably by cytokinins, and chromatin level regulation of WUS expression are also important in meristem size control, and can lead to bigger meristems when disrupted (Chickarmane et al. 2012; Gordon et al. 2009; Gruel et al. 2016; Kaya et al. 2001; Landrein et al. 2014). Here we report a new mutant with large meristems, *oki1*, that encodes an aspartyl tRNA synthetase. *Arabidopsis* encodes three aspartyl tRNA synthetase homologs, with two predicted to encode cytoplasmic proteins, and OKI1 is the only to encode a protein that is predicted to be localized to organelles. We found using a functional YPet fusion that the OKI1 protein product localizes predominantly to mitochondria. Not surprisingly, *oki1* null mutants were lethal, however we identified a viable, and therefore weak, allele uncovering a function of the *OKI1* aspartyl tRNA synthetase in meristem maintenance that could not be elucidated from null alleles.

Analysis of meristem structure in the *oki1* weak allele provided important insights into meristem control. First, the organization of cells into regular clonal cell layers was disrupted in both shoot and root, however the meistems were able to function and complete the plant lifecycle, including production of viable flowers and seeds. This supports the idea that plant development is controled by positional information, and even where clonal divisions are observed, such as in meristems, such regular divisions are not essential for maintaining stem cell niches (Smith et al. 1996). Moreover, we found that expression of both CLV3 and WUS genes were maintained as correctly positioned separate domains, further supporting the idea that these domains are established using positional cues rather than a dependency on cell lineage. In addition to the meristems, leaf development was also severely disrupted in *oki1* mutants; the first leaves produced were narrow and strap shaped, and later leaves were near normal width but asymmetrically lobed. Therefore full *OKI1* function appears necessary for many aspects of *Arabidopsis* development.

What is the mechanism of *OKI1* function? AaRSs perform housekeeping roles in protein translation, by catalyzing the ligation of amino acids and their cognate transfer tRNAs to prepare substrates for protein translation (Vargas-Rodriguez et al. 2018). In plants, translation occurs in three different cellular compartments, the chloroplasts, mitochondria and cytosol (Berg et al. 2005). Translation in each of these compartments is necessary, as elimination of some chloroplast, mitochondria or cytosol aaRSs in *Arabidopsis* leads to embryo lethality, ovule abortion or gametophytic lethality (Berg et al. 2005). Null mutants in *OKI1* are also lethal, but the weak *oki1* allele is viable, with severe effects on meristem and organ growth. The *oki1* weak mutant phenotype differed from most meristem mutations, in that the cellular organization was highly irregular, and suggest that it may function through a different mechanism compared to the canonical meristem pathways. This weak allele has a single amino acid substitution in the conserved oligonucleotide-binding (OB) fold domain, which is involved in recognition of the anti-codon in the mRNA by the charged tRNA (Mirande 2017). Therefore this mutation is predicted to disrupt translation, presumably in mitochondria. Therefore it may affect the energy balance of cells, and it is not surprising that the phenotype is evident in meristems and developing primordia, where energy demand for growth is high. However, it is surprising that the mutation leads to meristem enlargement, since most similar mutations do not. One other developmental mutant in an organellar targeted RNA synthetase gene, a glycyl tRNA synthetase, is a weak allele of *EMBRYO DEFECTIVE DEVELOPMENT1* (*EDD1*), that is lethal in null alleles (Moschopoulos et al. 2012). Weak *edd1* mutants enhance *asymmetric leaves1 (as1)* phenotypes, and affect genes involved in leaf dorsiventral polarity, though again the mechanism is unknown, and no meristem phenotype was reported or is evident in *edd1* mutants. Leaf development is also abnormal in *defective chloroplasts and leaves* (*dcl*) mutants in *Arabidopsis* and tomato; *DCL* encodes a plastid targeted protein that functions in ribosomal RNA processing, but again no effect on meristem size was reported (Bellaoui and Gruissem 2004; Bellaoui et al. 2003). Other mutations with developmental phenotypes affect the cytoplasmic translational machinery, including ones in the *PIGGYBACK* (*PGY*) genes in *Arabidopsis*; *pgy* mutants were first identified in a screen for leaf polarity mutants in *Arabidopsis*, and encode ribosomal large subunit proteins (Pinon et al. 2008). Mutants in a related gene also affects leaf development in rice, for example *rice minute-like1* (*rml1*) mutants are smaller with defective vascular patterning and narrow leaf blades, and may have auxin related defects, however again no effect on meristem size or organization were reported (Zheng et al. 2016). *rml1* mutants also have small panicles, suggesting that shoot meristem size is reduced, which may reflect defects in cell growth or proliferation expected when ribosomes are compromised. Alternatively, it is possible that the specific developmental phenotypes of ribosomal mutants reflects a true regulatory role in development (Byrne 2009). However all of the plant mutants affecting ribosomes or translational machinery have abmormal leaf development with no obvious defect in meristems, and no reported example of meristem enlargement or fasciation, suggesting the *oki1* phenotype is unique. One possible mechanism to explain *oki1* phenotypes is that the OKI1 protein has an additional function, distinct from its role in translation. In mammals, aaRSs have alternative functions, such as in transcriptional control, extracellular receptor-mediated signaling or in mammalian target of rapamycin (mTOR) signaling (Schimmel 2018). In *Arabidopsis*, an *OKI1* homolog, the AspRS IMPAIRED IN BABA-INDUCED IMMUNITY 1 (IBI1) functions in a noncanonical way in plant defense to perceive β-aminobutyric acid (BABA), a nonprotein amino acid protecting plants against broad-spectrum diseases (Luna et al. 2014). BABA binds to IBI1 and blocks its L-Asparate binding site, switching the AspRS canonical activity of IBI1 to the noncanonical defense activity upon pathogen infection. However this is unlikely to be the case for *OKI1*, because unlike *ibi1* mutants, *oki1* mutants were not hypersensitive to BABA (Fig. S8). Another hypothesis is that disruption in mitochondrial translation could create a redox unbalance, common in mutations that affect mitochondrial function (Mignolet-Spruyt et al. 2016). Recently, roles for redox signaling in shoot and root meristem size control have been discovered. For example, histological staining found different types of reactive oxygen species (ROS) enriched in different shoot meristem domains; superoxide is enriched in the stem cells and promotes WUS expression, and differentiation is promoted in the peripheral zone by hydrogen peroxide. The function of these ROS species is illustrated by different mutants affecting ROS status, which have shoot meristem size defects (Zeng et al. 2017). Similar findings have been reported in rice, where a glutamyl-tRNA synthetase expressed in meristematic cell layers during anther development maintains cellular organization and regulates the population of male germ cells through the control of protein synthesis, metabolic homeostasis and redox status (Yang et al. 2018). The size of root meristems is also controlled by a redox mechanism, for example the UPBEAT1 transcription factor controls the balance between cell proliferation and differentiation, by controlling expression of peroxidase gene targets (Tsukagoshi et al. 2010). Redox control of meristem size in maize is also evident, as the glutaredoxin enzyme MALE STERILE CONVERTED ANTHER1 (MSCA1) controls activity of the FASCIATED EAR4 (FEA4) transcription factor (Yang et al. 2015). The *Aberrant phyllotaxy2* (*Abph2*) mutant has bigger meristems and is caused by dominant mutations in *MSCA1*, and *msca1* loss of function mutants have smaller shoot meristems (Yang et al. 2015). Redox signaling can also control shoot meristem size by modulation of plasmodesmata, for example severe changes in redox state in the *Arabidopsis gfp aberrant trafficking1* (*gat1*) mutant of *Arabidopsis* lead to a reduction in root meristem size and premature shoot meristem termination, presumably because of excessive callose deposition, potentially impacting the flow of nutrients and developmental signals (Benitez-Alfonso et al. 2009). Further study should elucidate the precise mechanism of *OKI1* in redox inbalance and its role in meristem size control.

## Materials and Methods

### Plant materials and growth conditions

Mutagenesis in *Arabidopsis* Col-0 was performed as described previously (Xu et al. 2011). The M1 progeny were allowed to self-fertilize, and mutants screened in the M2 population. The following lines were obtained from the Arabidopsis Biological Resource Center: *clv3-2* (Ler ecotype) (Fletcher et al. 1999), SAIL_358_B08 and the double transgenic line expressing *CLV3*∷*mGFP5-ER* and *WUS*∷*dsRED-N7* (Ler ecotype) (Gordon et al. 2007). *wus* mutant line (Ler ecotype) was kindly provided from Dr. Yuval Eshed (Weizmann Institute of Science, Israel). This allele in which a T-DNA is inserted into *wus* was found as part of an enhancer trap population screen (Eshed et al. 1999). *ibi1-1* mutant line (Col-0 ecotype) was kindly provided from Dr. Jurriaan Ton (The University of Sheffield, UK). WUS∷GUS line was kindly provided by Dr. Damianos Skopelitis (Cold Spring Harbor Laboratory, USA) (Skopelitis et al. 2018).

For all experiments, Col-0 ecotype was used as a control line or WT. All plants were grown on soil or Murashige and Skoog (MS) agar plates (Weigel and Glazebrook 2002) at 23°C under long-day (LD, 16 h light/8 h dark) conditions. *Arabidopsis* seeds were stratified on soil or MS plates in the dark at 4°C for 48-72 h before transferring them to growth conditions.

Transgenic *Arabidopsis* was obtained by *Agrobacterium*-mediated floral dip (Alonso and Stepanova 2014; Weigel and Glazebrook 2002). *wus oki1*, *clv3-2 oki1* and *SAIL_358_B08 oki1* double mutants, and *CLV3*∷*mGFP5-ER WUS*∷*dsRED-N7* in *oki1* lines were generated through genetic crosses, and identified in the F2 segregating population. *oki1*, *clv3-2* and SAIL_538_B08 were validated based on PCR genotyping. *wus* was identified by Basta screening and PCR genotyping, and *CLV3*∷*mGFP5-ER WUS*∷*dsRED-N7* were validated by GFP or dsRED detection by fluorescence microscopy. Primers used for genotyping are listed in Table S1.

### Gene mapping

The *oki1* mutant (Col-0 ecotype) was crossed to the Ler ecotype. The F1 progeny were allowed to self-fertilize, and in the F2 population *oki1* phenotype (Fig. 1B) segregated 3:1. DNA was collected from pooled *oki1*- and WT-like plants. Library preparation was carried out according to manufacturers instructions (NEB Next Ultra DNA Library Prep Kit for Illumina, New England BioLabs Inc) and paired end sequencing was performed on the Illumina platform at the Cold Spring Harbor laboratory (New York, USA). Sequencing data was analyzed with the short read analysis pipeline SHOREmap (Schneeberger et al. 2009). Reads were aligned to the *Arabidopsis* Col-0 reference genome (TAIR 10). SNPs detected by sequencing were converted to CAPS or dCAPS markers, and a final mapping interval supported by several recombinants on each side was defined by markers at 15.65 Mb and 16.34 Mb in the Col-0 reference genome.

### Molecular biology

Recombineering lines containing OKI1 fused to three copies of YPet (a YFP variant) (Nguyen and Daugherty 2005) at the C terminus (OKI1-3xYPet) were generated as described (Zhou et al. 2011). The JAtY59F05 clone and the recombineering cassette with 3xYPet (3xAraYPet-FRT-Amp-FRT) were kindly provided by Dr. Jose Alonso (North Carolina State University, USA). The Escherichia coli recombineering strain SW105 was from the National Cancer Institute (Maryland, USA). The cassette was introduced to the C terminus of OKI1 by PCR using primers 1 and 2 (Table S1). The 3xYPet-tagged JAtY59F05 clone was transformed into *Agrobacterium tumefaciens* (GV3101), and transgenic recombineering strains were selected by the procedure of Zhou et al. (2011) using the primers 3 and 4 (Table S1). The recombineering transgenic plants were screened by spraying Basta on soil.

### Shoot apical meristem (SAM) measurement

SAM measurement was performed as described previously (Balkunde et al. 2017). Briefly, *Arabidopsis* seedlings were harvested at 8 or 12 DAP under LD conditions and then fixed overnight in ice-cold FAA (10% formalin, 45% ethanol and 5% acetic acid) followed by dehydration through an ethanol series, and cleared with methyl salicylate (Sigma-Aldrich). Meristems were observed using Nomarski optics. The width and height of the meristems were measured and quantified using ImageJ-Fiji (Schindelin et al. 2012).

### Chemical staining

For Eosin Y staining of cells in the SAM, 10 DAP seedlings of WT and *oki1* were fixed in FAA (5% formalin, 5% glacial acetic acid, 45% ethanol) overnight at 4°C followed by dehydration through a 50-100% ethanol series. During the dehydration step, tissues were stained with 0.1% Eosin Y (Sigma-Aldrich) in 100% ethanol, and embedded into paraffin (PARAPLAST X-TRA; McCormick SCIENTIFIC). Sections (10 μm) were prepared using a microtome. For propidium iodide (PI) staining of cell wall in the root, the roots of WT and *oki1* were stained in 10 μg/ml PI for 5 minutes, rinsed and mounted in water.

For mitochondrial staining, protoplasts were isolated from rosette leaves of 2-week old transformants that expressed OKI1∷3xYPet driven under *OKI1* native promoter in *oki1* background using an enzyme solution (400 mM mannitol, 20 mM MES, 20 mM KCl, 10 mM CaCl2, 10 μg/ml BSA, 15 mg/ml cellulase [PhytoTechnology Laboratories], 3 mg/ml pectolyase [Sigma-Aldrich]). Isolated protoplasts were suspended in buffer (400 mM mannitol, 20 mM MES, 20 mM KCl, 10 mM CaCl2, 10 μg/ml BSA) containing 100 nM MitoTracker Red CMXRos (Molecular Probes) for mitochondrial staining. For staining in the SAM, shoot apices were embedded in 6% agar blocks, and sections were obtained with a vibratome and stained with MitoTracker Red CMXRos in PBS for 30 min.

For staining of ROS, superoxide anions and hydrogen peroxide, nitroblue tetrazolium (NBT) (Sigma-Aldrich) and 3,3’-diaminobenzidine (DAB) (Sigma-Aldrich) were used as described previously (Zeng et al. 2017). Briefly, 12 DAP seedlings were infiltrated with 1/2 liquid MS and either 1 mg/ml NBT and 50 mM potassium dihydrogen phosphate (pH 7.6) (for superoxide anion detection) or 1 mg/ml DAB and 10 mM disodium hydrogen phosphate (pH 6.5) (for hydrogen peroxide detection), and incubated in the dark for 10-20 h at room temperature. Stained plants were transferred into boiling ethanol/glycerin/glacial acetic acid solution (3:1:1) to terminate the staining, then fixed with paraformaldehyde (PFA) solution (2% paraformaldehyde, 0.1% DMSO in PBS) and embedded into 6% agar blocks for sectioning with a vibratome.

### Microscopy

Seedling images were taken with Nikon SMZ1500 (Nikon Instrument Inc) microscope to manually capture Z series, which were then merged using NIS elements to create focused images. Confocal images were obtained on a ZEISS LSM710 or LSM 780. For Eosin Y, 514 nm laser excitation and 538–680 nm emission spectra, for PI, 514 nm excitation and 566-718 nm emission spectra, for MitoTracker Red, 561 nm excitation and 572-621 nm emission spectra, for YPet, 514 nm excitation and 519-588 nm emission spectra, for GFP, 488 nm excitation and 493-541 nm emission spectra, for dsRED, 594 nm excitation and 599-641 nm emission spectra, for chloroplast autofluorescence, 633 nm excitation and 647-721 nm emission spectra were used. LSM files from the confocal were processed using ImageJ-Fiji (Schindelin et al. 2012).

### Multiple sequence alignment and construction of phylogenetic tree

For multiple alignment of amino acid sequences of AspRSs from eukaryotes (Fig. S1), amino acid sequences that possess high similarity with OKI1 were obtained from National Center for Biotechnology Information (NCBI) database (https://www.ncbi.nlm.nih.gov) by BLAST search. AspRS amino acid sequences from *Arabidopsis thaliana* (OKI1, TAIR ID: AT4G33760), *Medicago truncatula* (NCBI accession number: XP_003609716), *Populus trichocarpa* (NCBI accession number: XP_024463857), *Oryza sativa* (NCBI accession number: XP_015622473), *Brachypodium distachyon* (NCBI accession number: XP_003568687), *Physcomitrella patens* (NCBI accession number: XP_02435843), *Homo sapiens* (NCBI accession number: 4AH6_A), *Drosophila melanogaster* (NCBI accession number: NP_724018), *Saccharomyces cerevisiae* (NCBI accession number: PTN17328), *Chlamydomonas reinhardtii* (NCBI accession number: XP_001694949) were used. Multiple sequence alignment was performed by CLUASTAL OMEGA (https://www.ebi.ac.uk/Tools/msa/clustalo/) (Sievers et al. 2011). For constructing a phylogenetic tree for *Arabidopsis* aaRSs (Fig. S4), seven complete amino acid sequences that possess high similarity with OKI1 (AT5g56680, AT1G70980, AT4G17300, AT4G26870, AT4G31180, AT3G13490 and AT3G11710) were obtained from Phytozome v12.1 database (https://phytozome.jgi.doe.gov/pz/portal.html) by BLAST search. Multiple sequence alignment was performed by CLASTALW (https://www.genome.jp/tools-bin/clustalw) (Larkin et al. 2007). Neighbor-joining (NJ) tree was constructed by MEGA 7 (Kumar et al. 2016) with 1000 bootstrap.

### GUS staining

Seedlings were transferred to tissue culture plates containing GUS staining solution (50 mM Na-phosphate at pH 7.0, 10 mM EDTA, 0.1% triton X-100, 1 mg/ml of X-Gluc [5-bromo-4-chloro-3-indolyl-beta-D-glucuronic acid, BIOSYNTH], 5mM potassium ferricyanide and 5mM potassium ferrocyanide), placed under vacuum for 5 min, and then incubated in the dark at 37 ◦C overnight. Staining solution was removed, and tissues were cleared in 70% ethanol.

### Chemical treatment

Control line, *ibi1-1* and *oki1* seedlings were grown on MS medium plates for four days, and then transferred onto MS medium plates containing 0, 150, 500 or 1500 μM S/R-β-aminobutyric acid (Sigma-Aldrich). After ten days, the fresh weight of seedlings in each condition was measured.

### Statistical analysis

Data for multiple groups were analyzed by one-way analysis of variance with a post hoc multiple comparison test (Turkey’s HSD procedure) using R software (https://www.r-project.org).

## Acknowledgements

The authors thank Dr. Jose Alonso (North Carolina State University, USA), Dr. Yuval Eshed (Weizmann Institute of Science, Israel), Dr. Jurriaan Ton (The University of Sheffield, UK) and Dr. Damianos Skopelitis (Cold Spring Harbor Laboratory, USA) for kindly providing reagents, and Dr. Molly Hammell (Cold Spring Harbor Laboratory, USA) for assistance with sequence data analysis. The authors are also grateful to Ms. Irene Liao for assistance with mutant cloning.

