## Supplementary material for "Control of Shoot Meristem Size by an Aminoacyl tRNA Synthetase, *OKI1*, in *Arabidopsis*": Fig. S1-8 and table S1

**Supplementary figures**

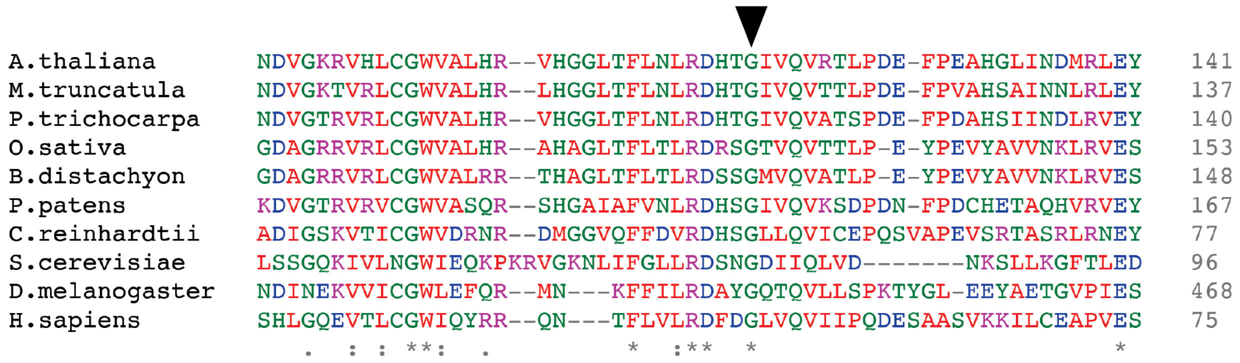


**Figure S1. Gly 116 in OKI1 is conserved among AspRSs in eukaryotes.** Alignment of amino acid sequence of AspRS from *Arabidopsis thaliana* (OKI1; AT4G33760), *Medicago truncatula* (XP_003609716), *Populus trichocarpa* (XP_024463857), *Oryza sativa* (XP_015622473), *Brachypodium distachyon* (XP_003568687), *Physcomitrella patens* (XP_02435843), *Chlamydomonas reinhardtii* (XP_001694949), *Saccharomyces cerevisiae* (PTN17328), *Drosophila melanogaster* (NP_724018) and *Homo sapiens* (4AH6_A). The Gly 116 that was replaced to Aspartic acid in *oki1* mutant is fully conserved in plants and animals (arrowhead).

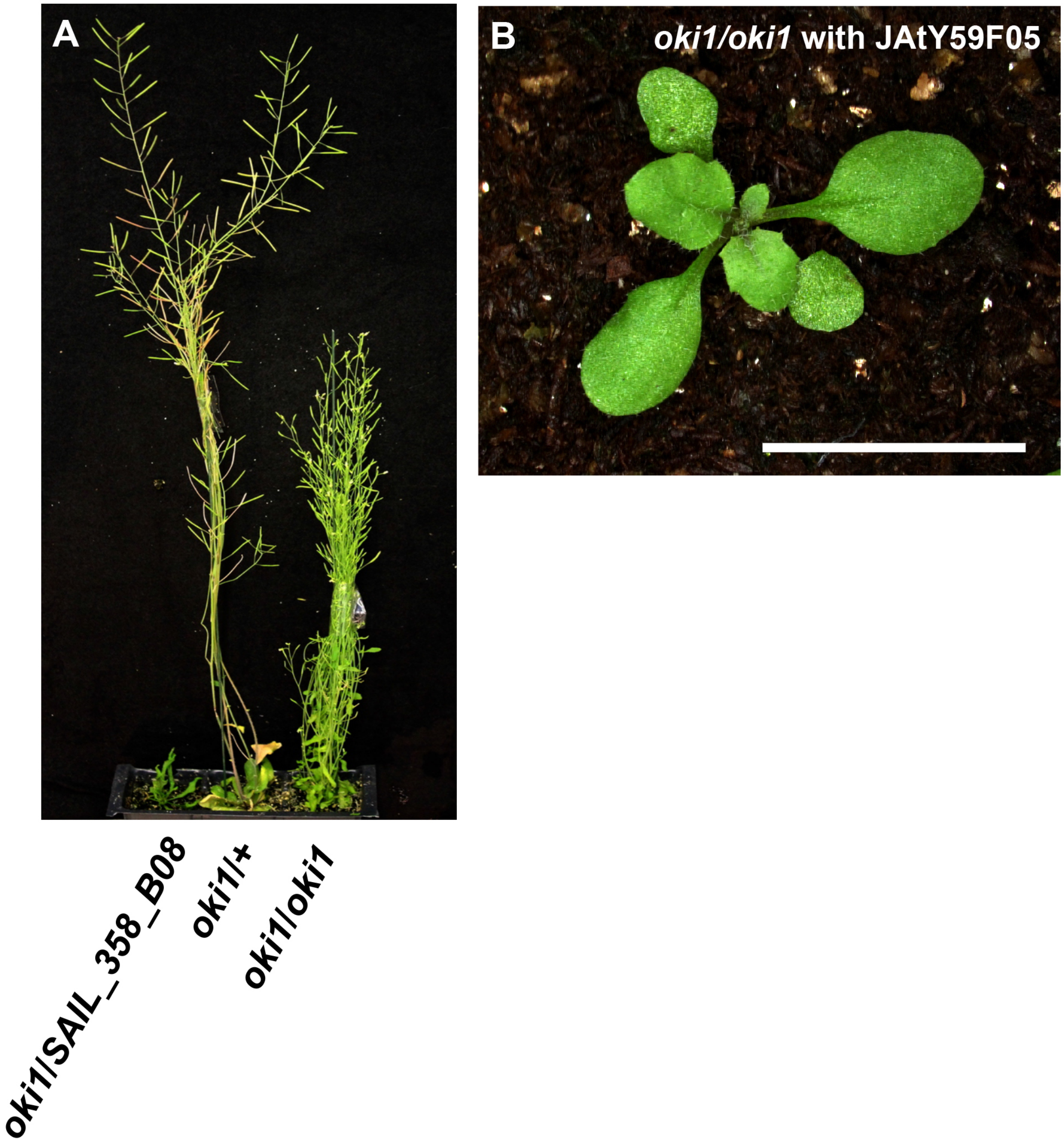


**Figure S2. Genetic complementation of *oki1*.** (A) *oki1/SAIL_358_B08* (weak allele / null) plants failed to complement *oki1*, and displayed enhanced growth defect phenotype of *oki1*. (B) Developmental phenotype of *oki1* (Fig. 1B) was complemented by introduction of TAC clone JAtY59F05 containing the *At4g33760* gene. Scale bar = 1 cm.

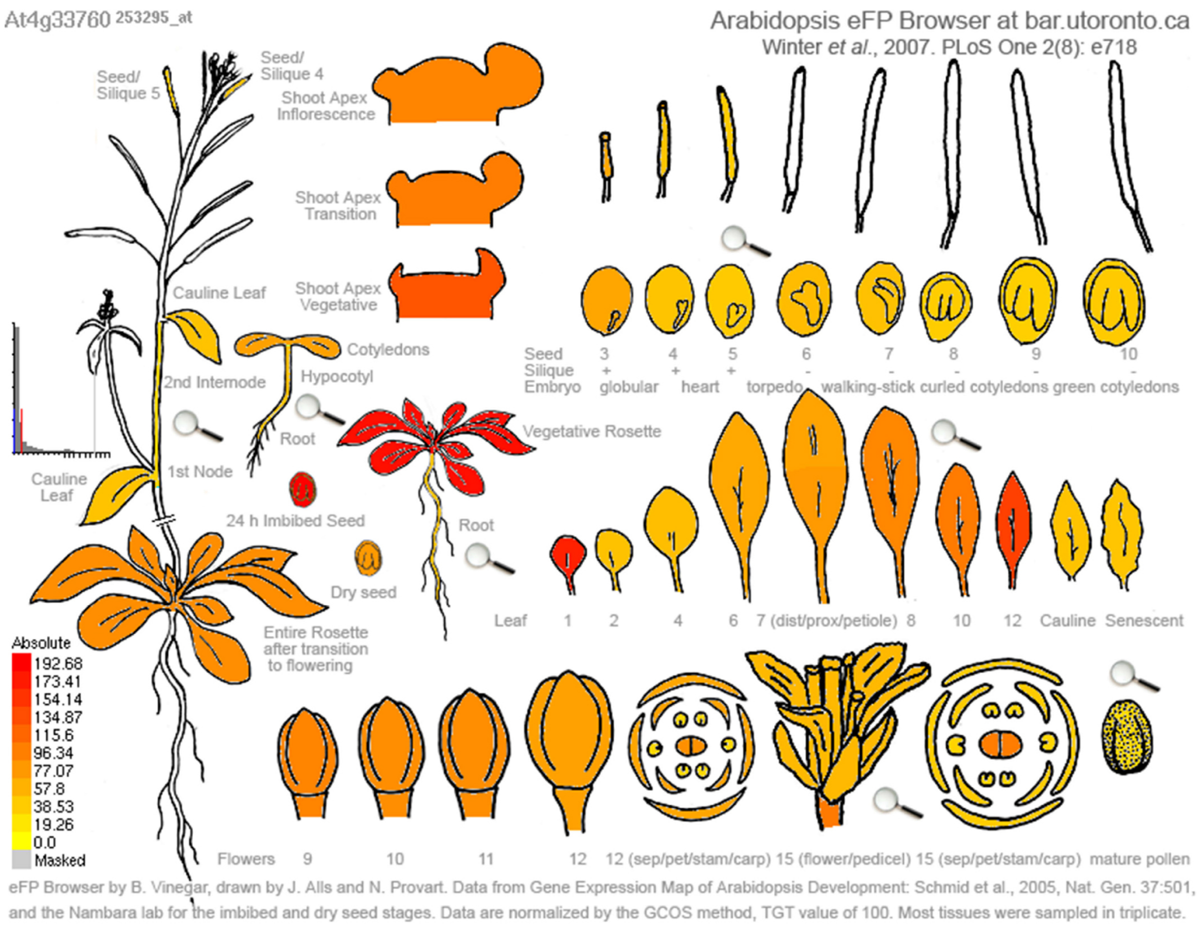


**Figure S3. *OKI1* expression in *Arabidopsis* development.** *OKI1* is expressed ubiquitously during *Arabidopsis* development. Figures were generated online using the eFP browser (http://bbc.botany.utoronto.ca/efp/cgi-bin/efpWeb.cgi). Relative expression levels of *OKI1* in various organs are shown via color scale, with red color indicating higher expression and yellow color indicating lower expression.

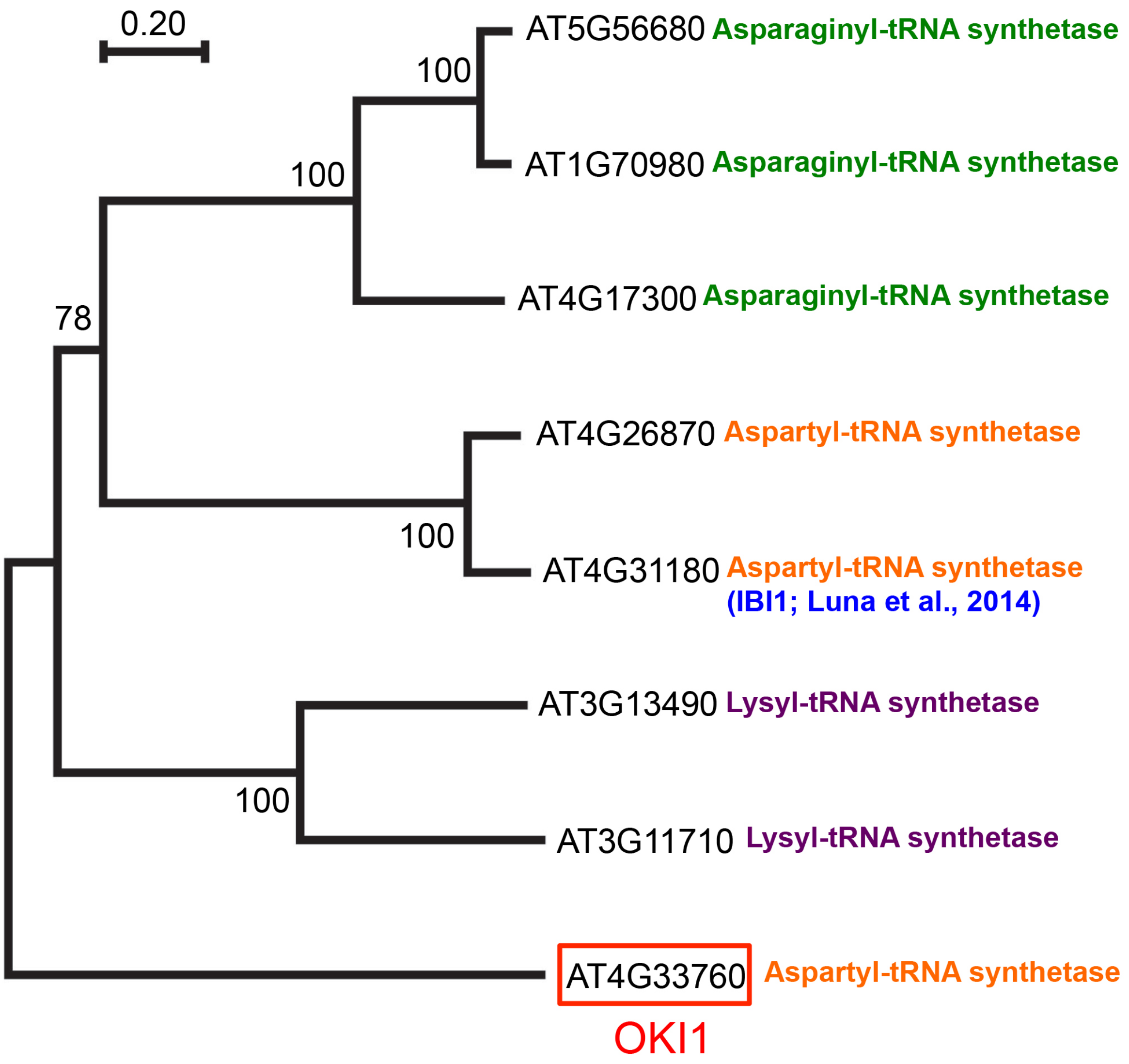


**Figure S4. *Arabidopsis* has three AspRSs.** A phylogenetic tree of 8 aminoacyl-tRNA synthetases from *Arabidopsis* conducted by MEGA7 software with the neighbor-joining (NJ) method for 1000 replicates bootstrap. In *Arabidopsis*, there are three AspRSs, AT4G33760 (OKI1), AT4G26870 and AT4G31180 (IBI1; Luna et al., 2014).

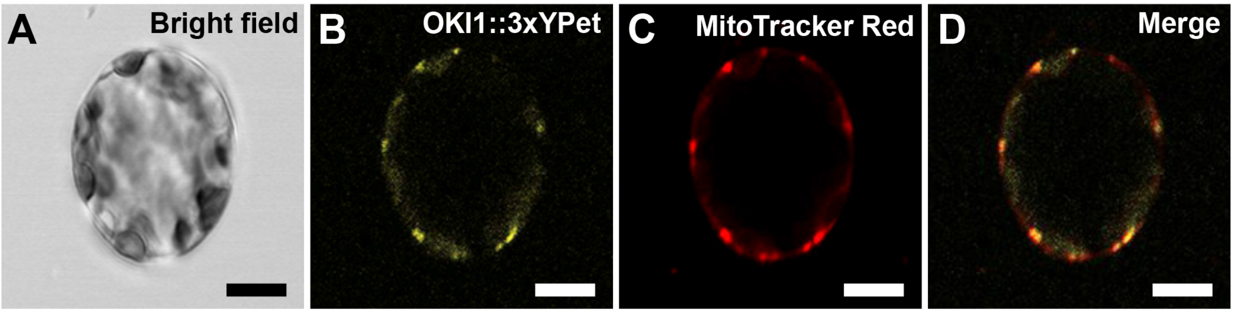


**Figure S5. Mitochondrial localization of OKI1 in protoplasts derived from leaf mesophyll cells.** (A) Bright field image of a protoplast derived from leaf mesophyll cells of the OKI1::3xYPet line driven by the native promoter in *oki1* background. (B) OKI1::3xYPet. (C) MitoTracker Red. (D) Merged image of B with C showed co-localization. Scale bar = 10 µm.

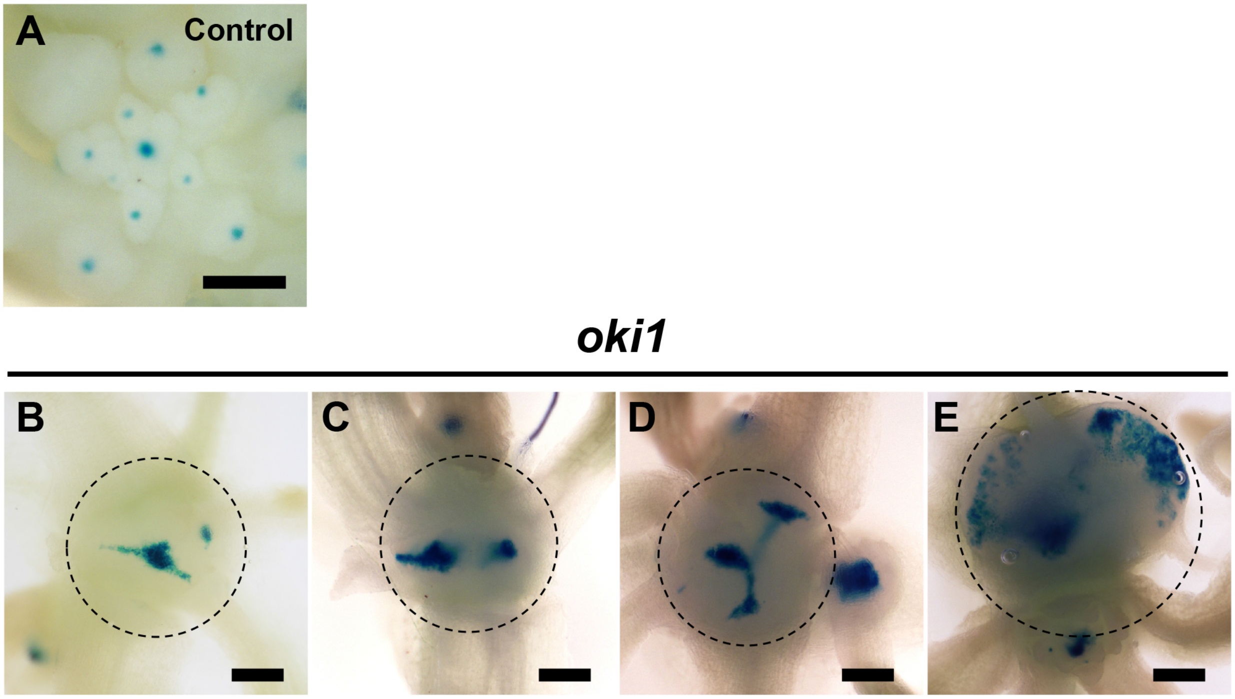


**Figure S6. Abnormal WUS expression in the SAM of *oki1* mutants.** (A-E) The expression patterns of promoter WUS fused GUS in control line (A) and *oki1* mutants (B-E). WUS expression site (blue region) was enlarged and/or split in the SAM of *oki1* mutants (blue regions in dashed circles). Scale bar = 200 μm.

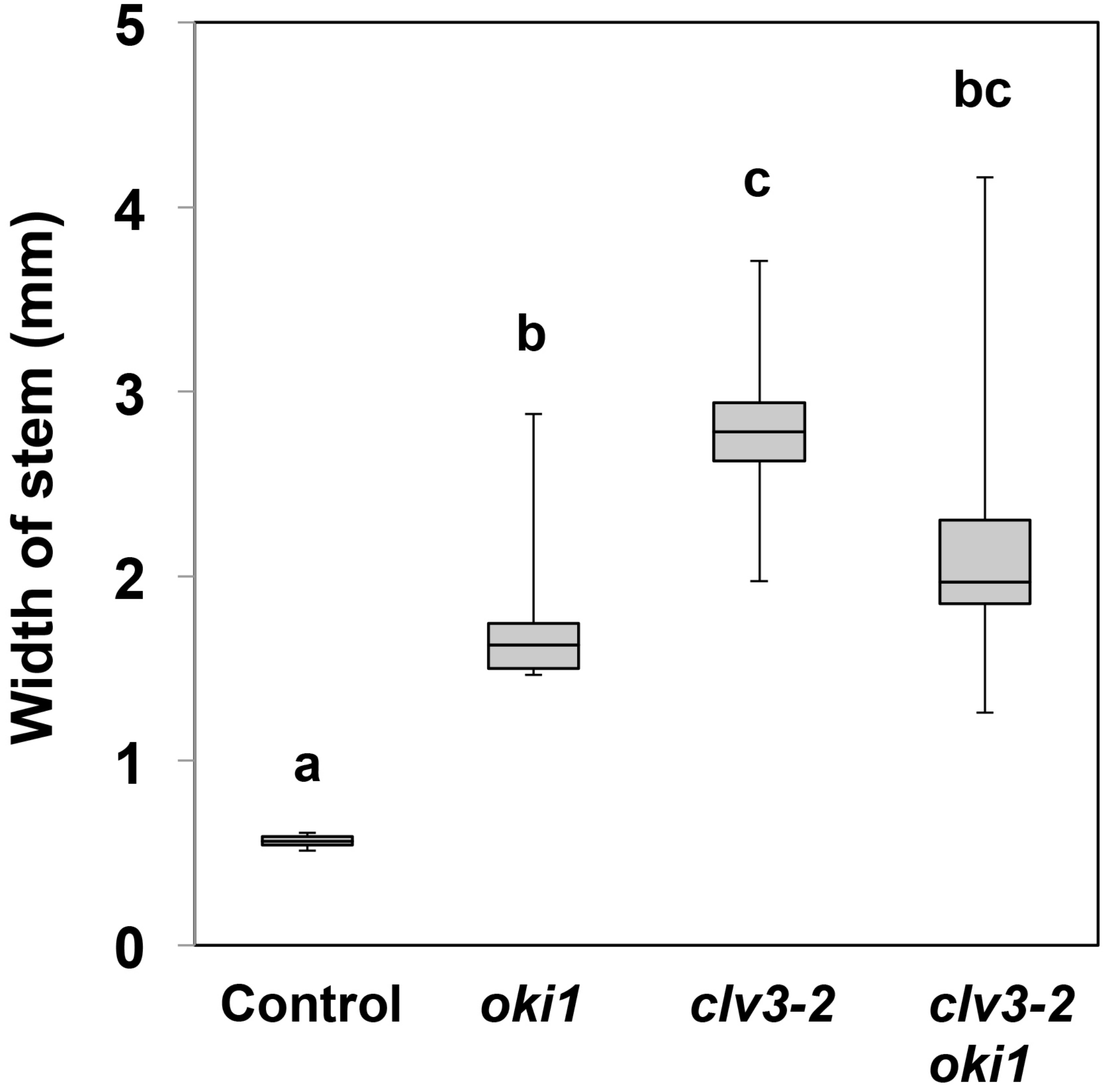


**Figure S7. *clv3* is epistatic to *oki1* in stem thickness control.** There was no significant difference in stem thickness between *oki1* or *clv3-2* single mutants and *clv3-2 oki1* double mutants, suggesting that *clv3-2* are epistatic to *oki1* in regulation of stem thickness. N = 7-10. Bars topped by different letters are significantly different at $P < 0.01$ (Tukey HSD test).

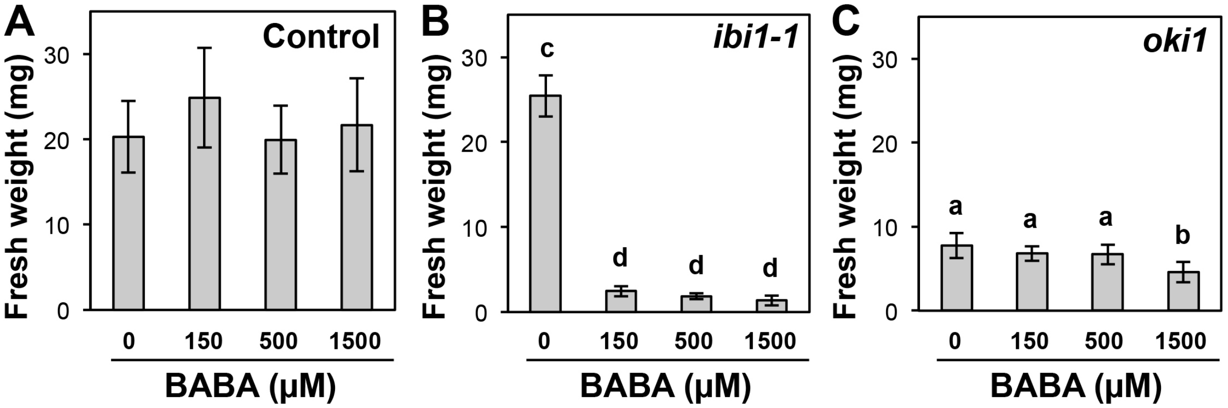


**Figure S8.** ***oki1* mutants are not hypersensitive to BABA.** (A, B) 150-1500 μM BABA did not affect to the growth of control line (A), but significantly inhibited the growth of *ibi1-1* mutants (B), as expected. (C) 150-500 μM BABA did not affect to the growth of *oki1* mutants, although it was decreased in presence of 1500 μM BABA. N = 8-10. Bars topped by different letters are significantly different at $P < 0.01$ (Tukey HSD test).

Table S1. Oligonucleotides used as primers in this study.

| Primer sets for genotyping. | | | |
|---|---|---|---|
| Mutants | Forward | Reverse | Note |
| *oki1* | GAGCTATGCGGCGAGTTATC | GAAAGTTCATTGCTGAGACGAA | PCR product was digested with HpaII. WT: 116bp / 113 bp, *oki1* : 229 bp. |
| *wus* | GGTCTTGCGAAGGATAGTGG | TTGCCCATCCTCCACCTACG | |
| *clv3-2* | CTCACTCAAGCTCATGCTCACG | GGGAGCTGAAAGTTGTTTCTTGG | Muller et al., 2008 |
| SAIL_358_B08 | CCTTATGATGCAGGCGAGAT | GCTGGCACTCTGAACAACAA | PCRs were performed with LBb1.3 primer (http://signal.salk.edu/tdnaprimers.2.html) |

| Primers for construction. | |
|---|---|
| Primer # | Sequence |
| 1 | CCTCGGAAGTCGATCCAAAGCAGCTTCAAGATCTCTCCATCCGCACCAAAGGAGGTGGAGGTGGAGCT |
| 2 | TGATGTTAAGAGTAAACAGAAGATACAATTGTTTGTGTTGAGAGCTATTAGGCCCCAGCGGCCGCAGCAGCACC |
| 3 | AAGATTGGTCAAGCATGGTTG |
| 4 | TGTCAAAAGTGGGAATTTTGC |
